# Targeting STING-ATM axis overcomes radiotherapy-induced immune suppression and restores anti-tumor immunity in nasopharyngeal carcinoma

**DOI:** 10.1101/2025.11.12.687945

**Authors:** Tzu-Tung Liu, Kai-Ping Chang, Yu-Nong Wang, Chun-Nan OuYang, Meng-Hsin Li, Ngan-Ming Tsang, Shih Sheng Jiang, Yun-Hua Sui, Chun-Mei Hu, Yung-Chih Chou, Hsuan-Chung Chen, Shu-Chen Liu

**Affiliations:** Department of Biomedical Sciences and Engineering, National Central University, Taiwan; Department of Otolaryngology-Head & Neck Surgery, Chang Gung Memorial Hospital, Taiwan; Department of Radiation Oncology, Taipei Tzu Chi Hospital, Taiwan; Molecular Medicine Research Center, Chang Gung University, Taiwan; Department of Radiation Oncology, China Medical University Hsinchu Hospital, Taiwan; National Institute of Cancer Research, National Health Research Institutes, Taiwan; Genomics Research Center, Academia Sinica, Taiwan; Department of radiation oncology, New Taipei Municipal TuCheng Hospital, Taiwan

**Keywords:** radiotherapy, single-cell RNA sequencing, radioresistance, STING, ATM, MPO, immune suppression, nasopharyngeal carcinoma

## Abstract

**Background:** Radiotherapy (RT) is a standard treatment for nasopharyngeal carcinoma (NPC), however, RT-induced immune suppression and tumor radioresistance limit durable cancer control, with approximately 20% of patients developing recurrence or metastasis after treatment. We investigated the molecular effects of RT on the immune landscape of NPC and identified a key role of the STING–ATM–MPO axis in RT-induced immune suppression, which is associated with regulatory T cell (Treg) expansion and radioresistance.

**Methods:** We analyzed leukocyte counts from 653 NPC patients before treatment and at multiple time points (week 1–2, week 3–4, and week 5–6) after RT. Single-cell RNA sequencing (scRNA-seq) was performed on peripheral blood mononuclear cell (PBMC) samples from stage I NPC patients (T1N0M0) treated with RT alone. Blood samples were collected before and after 18 Gy RT (in 2 Gy fractions). A randomized six-arm syngeneic mouse tumor model and in vitro cell studies were used to investigate interactions among STING–ATM signaling, immune modulation, and tumor response.

**Results:** In NPC patients, RT suppressed cGAS–STING signaling in monocytes and dendritic cells but promoted immunosuppressive programs, including PD-1 signaling, checkpoint expression, regulatory T cell expansion, and upregulation of TGFβ1, IL-10, VEGF, PDGF, and MDSC-associated pathways. In a syngeneic mouse model, co-treatment with the STING agonist diABZI reversed these effects and improved tumor control. Bulk RNA sequencing confirmed that RT suppressed cGAS–STING and ATM signaling in both spleens and tumors. Combination treatment restored this signaling, enhanced neutrophil degranulation, and inhibited PD-1 signaling and immune checkpoints. Immunohistochemistry showed that diABZI increased ATM activation, MPO expression, and reduced Treg infiltration in spleens and tumors. ATM activation correlated positively with MPO and inversely with FOXP3 immunoreactivity, suggesting that STING–ATM–MPO activation counteracts Treg-mediated immune suppression. In vitro, diABZI activated STING–ATM signaling in PBMCs, THP-1 cells, and Jurkat T cells, while inhibiting NPC cell growth partly via CYLD upregulation.

**Conclusions:** These findings link RT-induced suppression to inhibition of cGAS–STING signaling. Activation of STING-ATM-MPO axis counteracts Treg-mediated immune suppression, supporting the therapeutic potential of STING agonists as radiosensitizers in NPC.

## Introduction

Nasopharyngeal carcinoma (NPC) is prevalent in Southeast Asia [1, 2], where it exhibits a type II Epstein Barr virus (EBV) latent infection with an expression pattern restricted to the nuclear antigen, EBNA1; latent membrane proteins, LMP1, LMP2A and LMP2B; EBV-encoded small nuclear RNAs (EBERs); and BamHI A transcripts [2]. Limiting the number and levels of expressed viral proteins is likely a key strategy used by EBV to evade host immune surveillance during latency. Radiotherapy (RT) is the standard treatment for NPC and is often combined with chemotherapy for locally advanced disease. Whereas RT effectively controls primary tumors, approximately 20% of patients develop recurrence or metastasis, with radioresistance being a major cause of treatment failure [3]. The variability of the RT response is influenced by factors such as hypoxia, cancer stem cell persistence, immune activation, and DNA repair capacity [4]. Moreover, RT induces chronic inflammation, which promotes immune evasion and tumor progression [5]. Cytokines such as tumor growth factor beta 1 (TGFβ1), tumor necrosis factor alpha (TNFα), interleukin (IL)-1, IL-6, IL-4, and IL-10 are secreted post-irradiation by macrophages, fibroblasts and tumor cells, contributing to an immunosuppressive tumor microenvironment (TME) [6]. Inflammatory signaling pathways, including NF-κB (nuclear factor kappa B) and STAT3 (signal transducer and activator of transcription 3), further reinforce radiation resistance [7–9]. However, how local immune niches influence treatment response and recurrence risk remains poorly understood.

Innate immune sensing pathways, particularly those recognizing cytosolic DNA, are central to RT-induced immune responses. cGAS–STING (cyclic GMP-AMP synthase–stimulator of interferon genes) signaling, Toll-like receptors (TLRs), and RIG-I-like receptors (RLRs) detect nucleic acids from damaged or dying cells, initiate type I interferon (IFN) signaling and enhance antigen presentation [10], leading to T cell priming. STING activation drives IFN production, MHC class I expression and cytotoxic T cell recruitment [11, 12], enabling transformation of immunologically “cold” tumors into T-cell–inflamed phenotypes [13–15]. The role of STING signaling in NPC is less characterized. Several studies have reported that EBV-encoded products such as LMP1, BPLF1 and BGLF5 can suppress TLR and STING activation [16]. EBV also activates TRIM29, a protein that degrades STING, resulting in decreased IFN production in EBV-infected cells [17]. Preclinical studies have shown that STING agonists reprogram tumor-associated macrophages toward an M1-like phenotype and improve responses to RT and immune checkpoint blockade (ICB) therapy [18, 19]. Despite these benefits, STING activation can also contribute to immune suppression. High-dose RT (>12–18 Gy) induces TREX1 (three prime repair exonuclease 1), which degrades cytosolic DNA and inhibits cGAS–STING activation, allowing immune escape [20]. Furthermore, STING activation can recruit myeloid-derived suppressor cells (MDSCs) via CCR2 (C-C motif chemokine receptor 2) and promote immune exhaustion [21, 22], sustaining an immunosuppressive TME. These opposing outcomes highlight the complex and context-dependent role of STING in RT responses and underscore the need for mechanistic insights into its dual effects.

In this study, we investigated RT-induced systemic immune alterations using single-cell RNA sequencing (scRNA-seq) of peripheral blood mononuclear cells (PBMCs) from stage I (T1N0M0) NPC patients treated with RT alone. This early-stage, treatment-uniform cohort allowed us to isolate RT-specific immune effects without the confounding influence of concurrent chemotherapy. Functional analyses revealed that RT enhanced DNA-damage sensing, IFN signaling and antigen presentation, but suppressed cGAS–STING signaling in innate immune cells. Notably, enhanced regulatory T cell (Treg) signatures – immune checkpoints, programmed cell death 1 (PD-1) signaling, and tumor-promoting factors (TGFβ, VEGF, PDGF, IL-10) – and MDSC signatures were identified in post-RT samples, supporting the emergence of a radioresistant, suppressive immune response. In experiments testing the use of the STING agonist diABZI as a therapeutic strategy to counteract RT-induced immune suppression in a syngeneic mouse tumor model, we demonstrated that concurrent activation of the STING–ATM–MPO axis significantly attenuated RT-induced immune suppression.

## Materials and Methods

### Clinical samples

Retrospective clinical records of patients diagnosed with nasopharyngeal carcinoma (NPC) were obtained and reviewed. A total of 653 eligible NPC patients were included in the complete blood count (CBC) analysis (online supplemental Table 1). All patients received primary treatment with either radiotherapy or concurrent chemoradiotherapy at Chang Gung Memorial Hospital between January 2005 and December 2013. Exclusion criteria included: a second primary cancer within three years before or after treatment, a radiotherapy equivalent dose in 2 Gy fractions (EQD2) of less than 66.0 Gy, age under 18 years, use of surgery as the primary treatment, treatment duration exceeding 70 days, and distant metastases before completion of primary treatment. CBCs were available for 533, 604, and 516 patients at two, four, and six weeks after the start of radiotherapy, respectively. Ethical approval was granted by the Institutional Review Board of Chang Gung Memorial Hospital, Taiwan (IRB number: 202000190B0). Due to the retrospective nature of the study, the requirement for informed consent was waived. Fresh whole blood samples from NPC patients were obtained from Linkou Chang Gung Memorial Hospital, Taiwan, for single-cell RNA sequencing. Paired blood samples (pre-treatment and post-18 Gy in 2 Gy fractions) were collected from stage I (T1N0M0) NPC patients who received radiotherapy as the sole treatment. In total, six paired blood samples (twelve total) were obtained for single-cell RNA sequencing. Two pairs were excluded due to unexpected chemotherapy administration. All samples were approved by the Institutional Review Board of Chang Gung Memorial Hospital (IRB number: 202101548B0). Written informed consent was received prior to participation.

### Sample preparation for single-cell RNA sequencing

Peripheral blood mononuclear cells (PBMCs) were isolated using density gradient centrifugation. Whole blood collected in a heparin-coated tube was diluted with 0.9% normal saline and carefully underlaid with 15 mL Ficoll-Paque gradient (Sigma Aldrich). The mixture was centrifuged at 400 ×g for 40 min. PBMCs were collected from the buffy coat layer and washed twice with PBS. Contaminating red blood cells (RBCs) were removed using RBC lysis buffer. The processed cells were then resuspended at the desired concentration. Cell viability was assessed using Calcein-AM/PI double staining. Only cell suspensions with viability greater than 90% proceeded to library construction. Library preparation was performed using the Chromium™ Single Cell 3’ v3.1 reagent kit (10x Genomics) following the manufacturer’s protocol. Briefly, single-cell suspensions were loaded onto a microfluidic chip and partitioned into single-cell gel beads-in-emulsion (GEMs) using the Chromium Controller instrument. Subsequent steps, including barcoding, reverse transcription, cDNA amplification, and fragmentation, were carried out to generate 3’ single-cell cDNA libraries.

### Pre-processing of single-cell RNA sequencing data

Libraries were sequenced on the NovaSeq 6000 System (Illumina). Sequencing data were processed using Cell Ranger pipelines (version 7.0.0; 10x Genomics). Briefly, raw binary base call (BCL) files were demultiplexed and converted to FASTQ format. Reads were mapped to the human genome reference (GRCh38), and gene expression matrices were generated for downstream analysis. The Seurat R package (version 5.1.0) was used for quality control, global-scaling normalization, principal component analysis (PCA), clustering, and differential expression (DE) analysis [23]. Cells expressing fewer than 200 unique genes or more than 15% mitochondrial reads were excluded before cell typing analysis. Doublets were filtered out using the DoubletFinder R package [24], with expected doublet rates determined based on the multiplet rate table provided by 10x Genomics. Additional doublets were manually identified and removed based on marker gene expression. In total, 7,633 cells were excluded during quality control.

### Identification of cell types

A shared nearest neighbor (SNN) graph-based clustering approach was used to identify distinct cell populations among the 72,720 qualified immune cells. Cell types were annotated based on the expression of established markers: monocytes (FCN1, VCAN, CD14, and S100A12), T cells (CD3D, CD3E, and CD6), NK cells (NCR1, NCAM1, and XCL2), B cells (MS4A1, CD79A, and BLK), dendritic cells (CD1C, FCER1A, and CLEC9A), mast cells (TPSAB1, MS4A2, and HDC), neutrophils (FCGR3B and CEACAM8), and platelets (PPBP, GP9, and PF4).

### Pathway and signature enrichment analyses

Signature scores were calculated using the UCell R package (version 2.6.2) [25]. Gene set variation analysis (GSVA) [26] was performed using the GSVA R package (version 1.50.5).. Hallmark gene sets and C2 curated gene sets (including KEGG and Reactome databases) from the Molecular Signatures Database (MSigDB) were used as reference datasets. The Mann–Whitney test and Student’s t-test were used to identify key signatures affected by radiotherapy.

### Pseudotime trajectory and cell-cell communication analyses

Pseudotime trajectory of T cells was performed using Monocle3 package (version 1.3.7) [27] with default parameters. NicheNet (version 2.1.5) [28] was used to predict ligand-target interactions among different immune cell types. Monocytes and mDCs were designated as ligand sources (sender cells), while T cells were considered target (receiver) cells. Genes expressed in more than 5% of each population were classified as “expressed genes” for ligand activity analysis. Differentially expressed genes (DEGs) associated with immunosuppressive signatures and upregulated in the post-RT group were defined as “genes of interest,” while all expressed genes served as “background genes.” Prioritized ligands were identified using the predict_ligand_activities function in NicheNet, based on the Pearson correlation test, and ranked by the area under the precision-recall curve (AUPR).

### Cell culture and in vitro diABZI stimulation

BM1 (NPC-derived cell line, provided by Professor Yu-Sun Chang, Chang Gung University, Taiwan) [29], Jurkat (clone E6-1, human acute T-cell leukemia cell line, purchased from the Bioresource Collection and Research Center, Taiwan), human PBMCs (Lonza Bioscience), and THP-1 (purchased from the Bioresource Collection and Research Center, Taiwan) were cultured in RPMI-1640 medium supplemented with 10% fetal bovine serum (FBS; Peak) and 1% penicillin-streptomycin (Corning). NPC-TW06 cells (provided by Professor Chin-Tarng Lin, National Taiwan University Hospital) [30] were cultured in DMEM supplemented with 10% FBS. All cell lines used in this study were authenticated using 16-core short tandem repeat (STR)-PCR profiling (analyzed by the Bioresource Collection and Research Center, Taiwan). THP-1 monocytes were differentiated into macrophages by treatment with 150 nM phorbol 12-myristate 13-acetate (PMA; Sigma, P8139) for 16 hours, followed by replacement with complete RPMI-1640 medium and incubation for an additional three days. The mouse MTCQ1 cell line (provided by Professors Wan-Chun Li and Shu-Chun Lin, National Yang Ming Chiao Tung University, Taiwan) [31] were infected with ready-to-use lentiviruses carrying the firefly luciferase 2 reporter gene and a puromycin resistance gene, followed by selection with puromycin (8 μg/mL) (Invitrogen). The selected luciferase-expressing MTCQ1 cells (MTCQ1_Luc2) were maintained in DMEM supplemented with 10% FBS (Peak) and puromycin (2 μg/mL) (Invitrogen), and were used for syngeneic mouse tumor experiments. All cells were maintained at 37°C in a humidified atmosphere with 5% CO₂. Cells were routinely tested for mycoplasma contamination using Hoechst 33342 (Invitrogen) DNA staining and a PCR-based mycoplasma detection kit (MedChemExpress, HY-K0552). For diABZI treatment, cells were cultured in complete medium for 24 hours, then incubated in medium containing 1.5% FBS for 16 hours before treatment with 0.3 or 1 μM diABZI (MedChemExpress, HY-112921B) for 3, 6, or 24 hours, either alone or in combination with cisplatin (1 μg/mL) or vehicle.

### Syngeneic mouse tumor model

To assess in vivo immune responses, C57BL/6 mice were used. Murine cancer cells (MTCQ1_Luc2) were subcutaneously injected into the right leg of each mouse. Tumor growth was monitored using an in vivo imaging system (IVIS) until tumors reached approximately 50 mm³. Tumor volume was calculated as 0.5× Length × Width^2^. Mice were randomly assigned to six groups (n = 7–8 per group): untreated, diABZI-treated, 6 MV photon irradiation, 6 MeV electron beam irradiation, diABZI combined with 6 MV photon irradiation, and diABZI combined with 6 MeV electron beam irradiation. diABZI was administered intraperitoneally (2 mg/kg) daily for two consecutive days (days 5 and 6). Fractionated radiotherapy with either 4 Gy 6 MV photon or 6 MeV electron beam irradiation (Trilogy, Varian Medical Systems) was applied to the tumor site on days 8, 16, and 21. Mice were sacrificed on day 22, and tumor tissues and spleens were immediately harvested for analysis. Spleen size measurement was determined using Image J software. All animal experiments were conducted in accordance with laboratory animal care guidelines and approved by the animal committees of National Central University (NCU-112-026) and Chang Gung University (CGU110-161).

### Bulk RNA sequencing of mouse tissues

Mouse spleens and tumor tissues were stored in RNAlater solution (Qiagen) immediately after dissection. Tissues were dissociated using 18-gauge needles, followed by total RNA extraction using the RNeasy Mini Kit (Qiagen). RNA purity and quantification were examined using SimpliNano™ Spectrophotometers (Biochrom, MA, USA). RNA degradation and integrity were assessed using Qsep 100 DNA/RNA Analyzer (BiOptic Inc., Taiwan). Libraries were prepared using KAPA mRNA HyperPrep Kit (KAPA Biosystems, Roche) following manufacturer’s instructions. The library quality was evaluated using the Qubit 2.0 Fluorometer (Thermo Scientific) and Agilent Bioanalyzer 2100 system. Paired-end sequencing (2 × 150 bp) was performed on a NovaSeq X platform (Illumina) with a target depth of ∼50 million reads per sample. Reads were aligned to the mouse reference genome (GRCm38) using HISAT 2.1.0 [32] with default parameters. Gene-level read counts were generated using featureCounts [33] with GENCODE annotation [34]. Differential expression analysis was conducted using the DESeq2 R package (version 1.42.1), with normalization performed using the default median-of-ratios method. Differentially expressed genes were defined by an adjusted p-value < 0.05 and absolute log2 fold change > 1 (Benjamini and Hochberg’s approach). These genes were used for downstream GSEA [35] and functional analysis between groups.

### Real-time analysis of cellular responses using the xCelligence system

NPC cells (3 × 10³ cells in 100 μl/well) were plated into E-plates and grown in complete medium for one day. The medium was then replaced with medium supplemented with 1.5% FBS, and cells were incubated overnight prior to diABZI stimulation. Cell impedance was recorded every 10 minutes for 5 days. Vehicle (DMSO) and various concentrations of diABZI (30 nM, 300 nM, 10 μM, and 50 μM) were added to the culture to evaluate the concentration–response curves of NPC cells. Cell growth values were normalized to the time point of diABZI stimulation as indicated. All experiments were performed in triplicate.

### Western blotting

Cells were lysed in NP-40 lysis buffer containing 1 mM PMSF, PhosSTOP™ Phosphatase Inhibitor (Roche, catalog No. 4906837001), and cOmplete™ Protease Inhibitor Cocktail (Roche, catalog No. 4693132001). Equal amounts of protein were separated by sodium dodecyl sulfate-polyacrylamide gel electrophoresis and transferred to polyvinylidene fluoride membranes. Blots were probed with primary antibodies against phospho-STING (Ser366) (Cell Signaling, 19781, 1:2000), STING (Cell Signaling, 13647, 1:2000), phospho-ATM (Ser1981) (Epitomics, 2452, 1:3000), ATM (Epitomics, 1549, 1:2000), phospho-H2AX (S139) (GeneTex, GTX636713, 1:4000), H2AX (GeneTex, GTX108272, 1:500–3000), phospho-IRF3 (S396) (Cell Signaling, 29047, 1:500), IRF3 (Cell Signaling, 11904, 1:500), MPO (Abcam, ab208670, 1:500–1000), PARP (GeneTex, GTX636805, 1:100,000), CASP1 (GeneTex, GTX637906, 1:8000–160,000), IL-1β (GeneTex, GTX636889, 1:3000), CYCD (GeneTex, GTX636662, 1:1000), phosphor-IκB (Ser32) (Abcam, 92700, 1:2000), IκB ((Santa Cruz Biotechnology, sc271, 1:2000), and GAPDH (Abcam, ab9484, 1:5000). Blots were then incubated with a horseradish peroxidase (HRP)-conjugated secondary antibody and developed using an enhanced chemiluminescence detection reagent (GE Healthcare).

### Immunohistochemistry (IHC)

IHC staining of mouse spleen and tumor sections was performed using the Novolink detection system (Leica) according to the manufacturer’s instructions. Briefly, tissue sections were deparaffinized and subjected to antigen retrieval in citrate buffer (pH 6.0) or EDTA buffer (pH 8.8) at 98 °C for 5 minutes. Endogenous peroxidase activity was blocked using 3-4% (v/v) hydrogen peroxide. After washing, non-specific protein binding was blocked with Novocastra Protein Block Buffer (Leica, RE7102). Tissue sections were then washed and incubated for 60 minutes with primary antibodies against phospho-ATM (Ser1981) (GeneTex, GTX30636, 1:500), MPO (Abcam, ab208670, 1:700), FOXP3 (Abcam, ab20034, 1:200), or TGFβ1 (Abcam, Ab215715, 1:400). This was followed by incubation with a secondary polymer (Leica, Novolink Polymer, RE7140-CE or ScyTek, PolyTek polymerized HRP kit) for 15 minutes and development with diaminobenzidine (DAB) for 7–10 minutes. For the negative control, the primary antibody was omitted and replaced with antibody dilution buffer containing an equivalent amount of IgG from non-immune rabbit or mouse serum. Stained sections were imaged using an Olympus IX83 microscope.

### Immunofluorescent double imaging in tissue sections

Immunofluorescent double staining was performed to evaluate the localization of different proteins in mouse spleen and tumor sections. Deparaffinization, rehydration, and antigen retrieval followed the same protocol as the IHC procedure. Tissue sections were permeabilized with 0.03% Triton X-100 in 3% BSA for 2 minutes and then blocked with normal goat serum blocking solution (BioGenex, HK112) for 30 minutes. Primary antibodies against phospho-ATM (Ser1981) (Epitomics 2452, 1:800), FOXP3 (Abcam, 1:200), TGFB1 (R&D Systems, MAB240, 1:50), CYCD (GeneTex, GTX636662, 1:700), or IRF (Cell Signaling, 11904, 1:800) were incubated overnight at 4°C. After washing, tissue sections were incubated with secondary antibodies conjugated to Alexa Fluor™ 594 or Alexa Fluor™ 488 (Invitrogen) for protein visualization. Hoechst 33342 was used for nuclear staining.

### Quantitative real-time reverse transcription-polymerase chain reaction (QRT-PCR)

Total RNA was extracted using the RNeasy Mini Kit (QIAGEN). cDNA synthesis was performed with the First Strand cDNA Synthesis Kit (Roche). Diluted cDNA was used for QRT-PCR with SYBR Green Supermix (BIO-RAD) on a LightCycler 96 system (Roche), following the manufacturer’s protocol. Primers used in this study are listed in Supplemental Table 5. Gene expression was normalized to ribosomal protein L13a (RPL13A). All assays were performed in at least triplicate.

### Statistical analysis

For single-cell transcriptomic data, DEGs among cell types or between the pretreatment and RT groups were identified using the FindAllMarkers and FindMarkers functions in the Seurat R toolkit [36], respectively. Statistical significance between groups in the scRNA-seq data was determined using the Mann-Whitney U test. GraphPad Prism 10 (GraphPad Software) was used to assess the significance of IHC immunoreactivity across groups using Fisher’s exact test. Spearman’s correlation analysis was performed to evaluate correlations between target protein expressions in tissue samples. The Mann-Whitney U test was used to compare differences among in vivo experimental groups. All statistical tests were two-sided, and P values < 0.05 were considered statistically significant.

### Data availability

All data supporting the findings of this study are available within this published article, its supplemental data files, and from the corresponding author upon reasonable request. The raw and processed single-cell data have been deposited in the Gene Expression Omnibus (GEO) under accession number GSE279565. The bulked RNA sequencing data of mouse spleens and tumor tissues have been deposited in GEO as well (GSE298449). The online version of the supplemental data will be available on the journal’s official website upon publication.

## Results

### RT reprograms immune landscape and function

We analyzed PBMCs from 653 NPC patients before treatment and at multiple time points (week 1–2, week 3–4, and week 5–6) post RT (online supplemental Table 1). Lymphocyte counts showed the most pronounced decline, decreasing by 47.84% (from 1191 to 621.2 cells/μl, *P* < 0.0001) by week 3–4 and by an additional 25.79% (from 621.2 to 461 cells/μl, *P* < 0.0001) by week 5–6 (figure 1A). Neutrophils and monocytes declined more gradually. Specifically, neutrophil counts decreased by 20.48% (from 4518 to 3593 cells/μl, *P* < 0.0001) in week 5–6 post RT (figure 1B), whereas monocyte counts decreased by 4.92% (from 457.4 to 434.9 cells/μl, *P* = 0.0135) in week 3–4 and by 16.83% (from 434.9 to 361.7 cells/μl, *P* < 0.0001) in week 5–6 post RT (figure 1C). Most NPC patients received concurrent chemoradiotherapy, which may contribute to the leukocyte decline shown in figure 1A–C.

**Figure 1.**
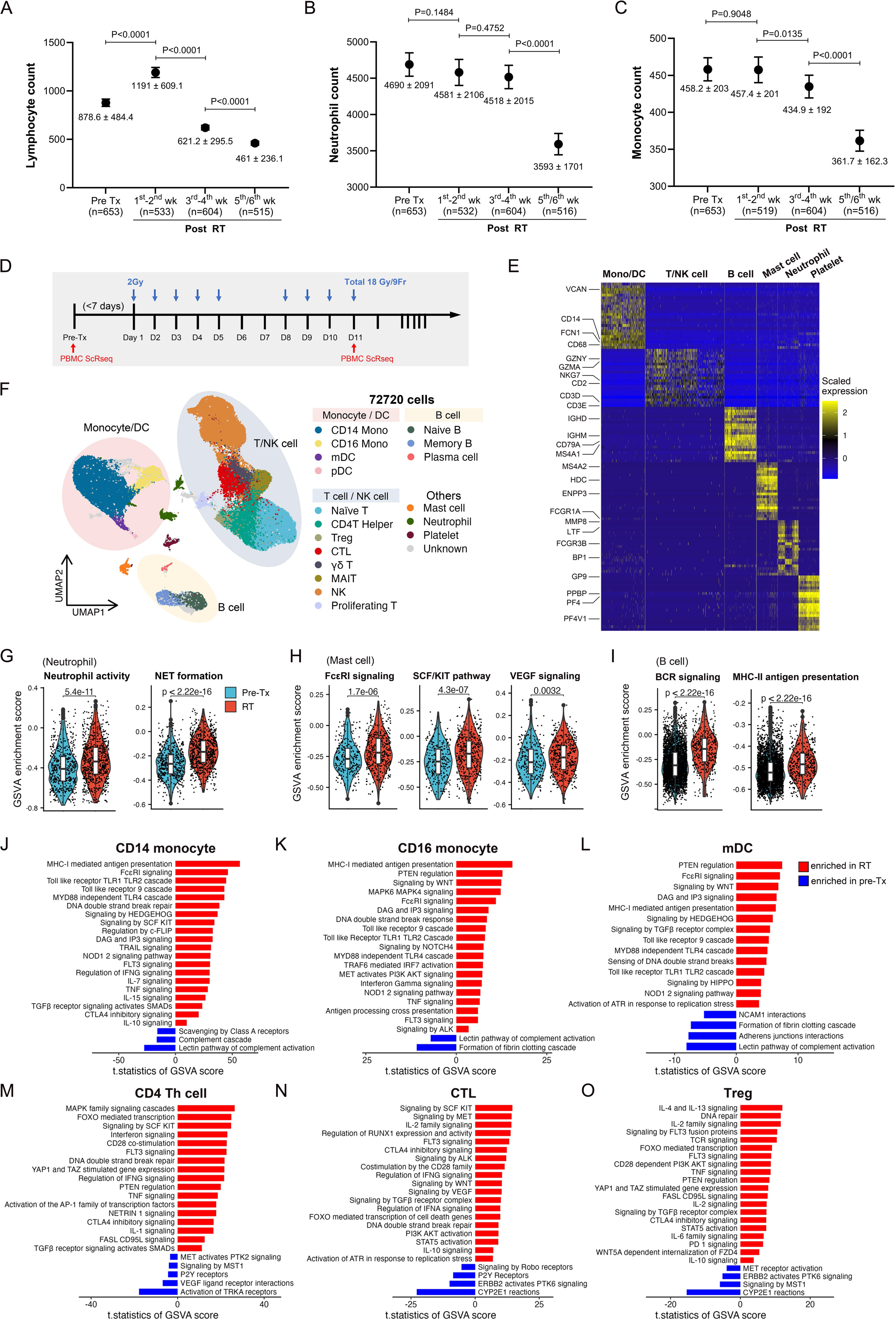
RT-induced alterations in systemic immune populations. (**A–C)** Leukocyte counts in PBMCs from NPC patients (n = 653) before treatment and during weeks 1–2, 3–4, and 5–6 post RT. (**A)** Lymphocytes; (**B)** neutrophils; (**C)** monocytes. Data are presented as means ± 95% confidence interval (paired *t* test). (**D)** Schematic diagram of clinical sample collection for single-cell transcriptomic analysis. (**E)** Expression heatmap of representative marker genes for distinct cell types. Color scale indicates expression levels from blue (low) to yellow (high). **(F)** UMAP plot of single-cell data grouped into annotated immune cell types. Each dot represents one cell. (**G)** GSVA enrichment scores for neutrophil activation and NET formation. (**H)** GSVA enrichment scores for FcεRI signaling, SCF/KIT signaling, and VEGF signaling in mast cells. **(I)** GSVA enrichment scores for BCR signaling and MHC-II antigen presentation in B cells. (G–I) Pre-treatment (Pre-Tx) and post-RT samples are shown in blue and red, respectively (Mann–Whitney U test). (**J–O)** GSVA enrichment scores for CD14^+^ monocytes (**J**), CD16^+^ monocytes (**K**), mDCs (**L**), CD4⁺ T helper cells (**M**), CTLs (**N**), and Tregs (**O**) comparing post-RT and pre-treatment samples. Enrichment scores (t-statistics) are shown in red (enhanced) and blue (reduced).

To isolate RT-specific immune effects, we performed scRNA-seq on PBMCs from stage I NPC patients (T1N0M0) who received RT alone. Blood samples were collected before and after 18 Gy RT (in 2-Gy fractions) (figure 1D), corresponding to the period before the steep lymphocyte decline. Libraries from six paired blood samples were generated and subjected to scRNA-seq. Two paired samples were excluded from analysis owing to unexpected chemotherapy administration. After quality control tests, we identified 72,720 immune cells across major immune populations, including monocytes, dendritic cells (DCs), T cells, NK cells, B cells, mast cells, and neutrophils. Marker gene mapping revealed distinct expression patterns for specific immune cell types (figure 1E). Uniform Manifold Approximation and Projection (UMAP) [37] was used to visualize cellular clusters (figure 1F). Representative marker genes included *CD3D* and *CD3E* for T cells; *CD68*, *FCN1* (ficolin 1) and *CD14* for myeloid cells; and *CD79A* and *MS4A1* (membrane-spanning 4-domains A1) for B cells.

A Gene Set Variation Analysis (GSVA) analysis of scRNA-seq results revealed that RT enhanced neutrophil activity and neutrophil extracellular trap (NET) formation (figure 1G and online supplemental Table 2); high-affinity IgE receptor (FcεRI) signaling, stem-cell factor (SCF)/KIT pathway activity, and vascular endothelial growth factor (VEGF) signaling in mast cells (figure 1H); and B-cell receptor (BCR) signaling and MHC-II antigen presentation in B cells (figure 1I). The GSVA results also highlighted differential pathway activities in CD14⁺ monocytes, CD16⁺ monocytes, myeloid dendritic cells (mDCs), CD4⁺ Th cells, cytotoxic lymphocytes (CTLs), and Tregs (figure 1J–O). RT increased antigen processing and presentation pathways, including class I MHC-mediated antigen processing and presentation, Toll-like receptor (TLR) signaling, FcεRI signaling, NOD2 (nucleotide binding oligomerization domain containing 2) signaling, and diacyl glycerol/inositol 1,4,5-trisphosphate (DAG/IP_3_) signaling in CD14⁺ monocytes, CD16⁺ monocytes, and mDCs (figure 1J–L). These pathways support antigen uptake, processing and presentation, and thereby enhance immune responses. Notably, T-cell subsets showed alterations in both activation (e.g., CD28 co-stimulation, IL-2 signaling, and TNF signaling) and suppression (e.g., PD-1, CTLA4, and IL-10 signaling) (figure 1M–O), indicating a dynamic immune homeostatic response following RT.

### RT enhances DNA-damage–sensing pathways but suppresses cGAS–STING signaling

Ionizing radiation induces the release of DNA or RNA fragments into the cytosol of damaged or dying cells, triggering a cascade of signaling events that initiate inflammation and innate immune responses. A GSVA revealed that RT increased the enrichment of cytosolic DNA sensor and DNA-repair pathway genes in innate and adaptive immune cells, except for plasma cells and mast cells (figure 2A and B). RT also altered the expression profiles of genes involved in antigen presentation and IFN signaling (figure 2C and D). Notably, several IFN-related genes, including *IFIT1* (interferon-induced protein with tetratricopeptide repeats 1), *IFIT3*, *ISG15* (ubiquitin-like modifier), *RSAD2* (radical S-adenosyl methionine domain-containing 2), and *MX1* (MX dynamin-like GTPase 1) were downregulated in monocytes and mDCs. A quantitative RT-PCR analysis of the same paired samples analyzed by scRNA-seq confirmed RT-induced downregulation of *IFNA* (interferon alpha complex region), *IFNB*, *IFIT1*, and *IFIT3* in PBMCs from NPC patients (figure 2E). Despite activating DNA damage sensing pathways, RT suppressed cGAS–STING signaling in monocytes and DCs (figure 2F). Key cGAS-STING pathway genes, including *CGAS* (cyclic GMP-AMP synthase), *STING1* (stimulator of interferon response cGAMP interactor 1), *CASP1* (caspase 1), *BAX* (BCL2-associated X), and *SURF4* (surfeit 4), were downregulated in monocyte and DC populations (figure 2G and H). A correlation analysis showed that cGAS–STING signaling was negatively correlated with *TGFB1* and *FOXP3* (forkhead box P3) expression in NPC tumor tissues (n = 113, GSE102349) (figure 2I), suggesting that suppression of the cGAS-STING pathway contributes to RT-induced immunosuppression. Together, these findings indicate that RT enhances DNA sensing while simultaneously suppressing cGAS-STING signaling, potentially promoting immune evasion.RT induces Treg expansion and enhances immunosuppressive pathways

**Figure 2.**
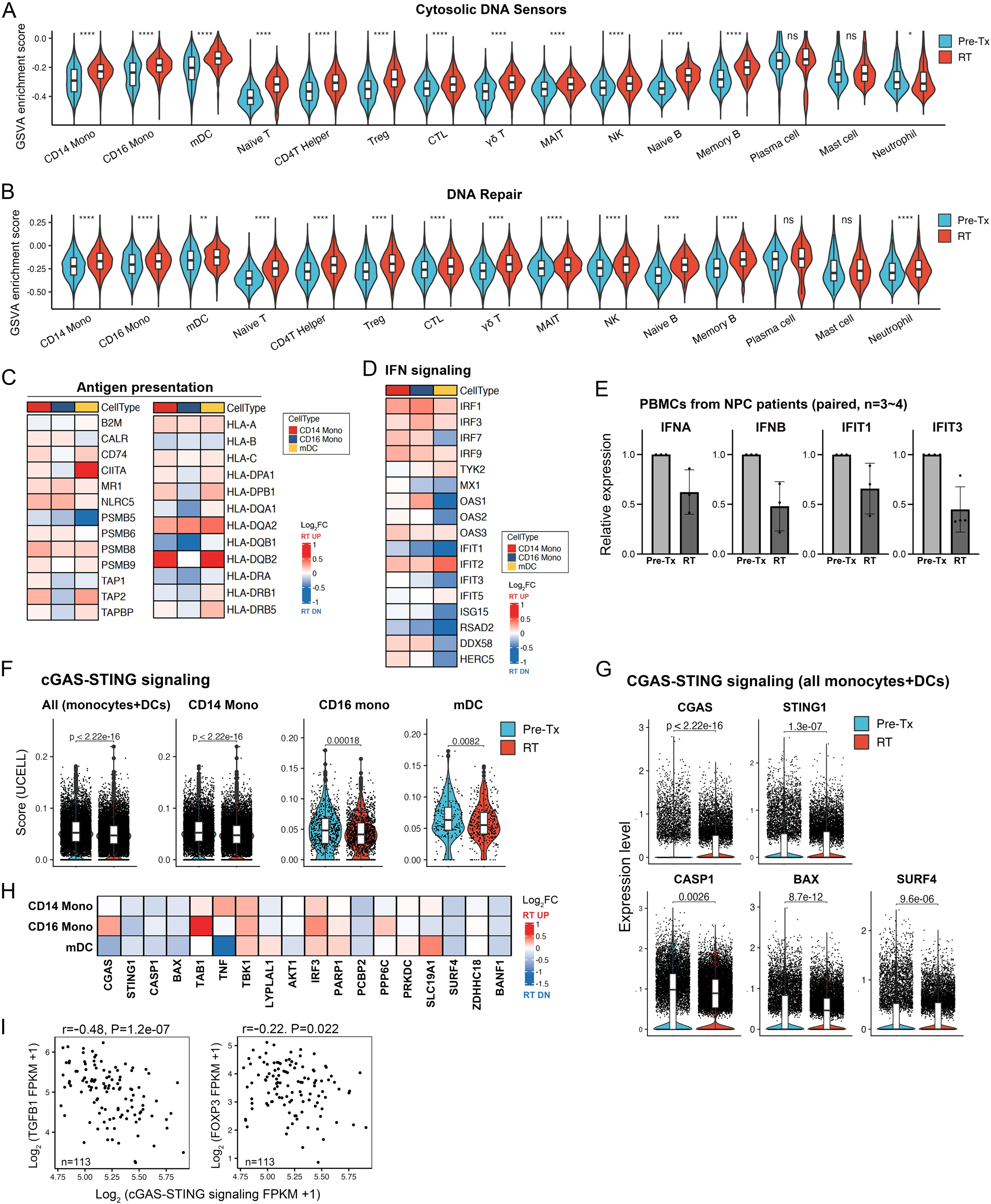
RT modulates DNA-sensing pathways in immune cells. **(A** and **B)** GSVA enrichment scores for cytosolic DNA sensors (**A**) and DNA repair (**B**) across innate and adaptive immune cells. (**C** and **D)** Heatmaps of genes involved in antigen presentation (**C**) and IFN signaling (**D**) in monocytes and mDCs. (**E)** Quantitative RT-PCR validation of *IFNA*, *IFNB*, *IFIT1* and *IFIT3* expression in paired pre-Tx and RT PBMCs from the same samples used in scRNA-seq. (**F)** Signature score for cGAS–STING signaling in CD14^+^ monocytes, CD16^+^ monocytes, and mDCs. Blue, pre-treatment (Pre-Tx) group; red, post-RT group. Each dot represents a single cell. Signature scores were calculated using the UCell R package (Mann-Whitney U test). (**G)** Expression levels of key genes in the cGAS–STING pathway across all monocyte and DC populations (Mann-Whitney test). (**H)** Heatmap of representative differentially expressed genes (DEGs) involved in cGAS–STING signaling in monocytes and mDCs. Expression values are presented as log2-transformed fold changes (Log_2_FC) of RT versus Pre-Tx. Red, RT-upregulated genes; blue, RT-downregulated genes. (**I)** Correlation analysis between cGAS–STING signaling scores and the expression of *TGFΒ1* (left) and *FOXP3* (right) in NPC tumor tissues (Spearman’s correlation, data from GSE102349).

A further analysis of RT-induced functional changes in T cells showed that the proportion of naïve T cells was markedly reduced post RT (figure 3A). RT also downregulated the expression of the naïve T cell markers, *SELL* (selectin L), *CCR7* (C-C motif chemokine receptor 7), *LEF1* (lymphoid enhancer binding factor 1) and *TCF7* (transcription factor 7, T cell specific), in both CD4⁺ and CD8⁺ T cells (figure 3B). Other subsets, including Tregs, CTLs, γδT cells, MAIT (mucosal-associated invariant T) cells, proliferating T cells and NK cells, remained relatively stable or increased. scRNA-seq analysis revealed a marked expansion of CD4⁺ T cells expressing *FOXP3*, *TGFΒ1*, *IL2RA* and multiple immune checkpoint molecules, indicating an increase in immunosuppressive T-cell subsets following RT (figure 3C). Within the Treg population, RT significantly elevated the expression of *FOXP3*, *TGFΒ1*, *IL2RA* and *CTLA4* (cytotoxic T-lymphocyte–associated protein 4), which are central to Treg identity and suppressive function (figure 3D). However, other exhaustion-associated markers, including *PDCD1* (programmed cell death 1), *LAG3* (lymphocyte activating 3), *HAVCR2* (hepatitis A virus cellular receptor 2), and *TIGIT* (T cell immunoreceptor with Ig and ITIM domains), showed no significant changes.

**Figure 3.**
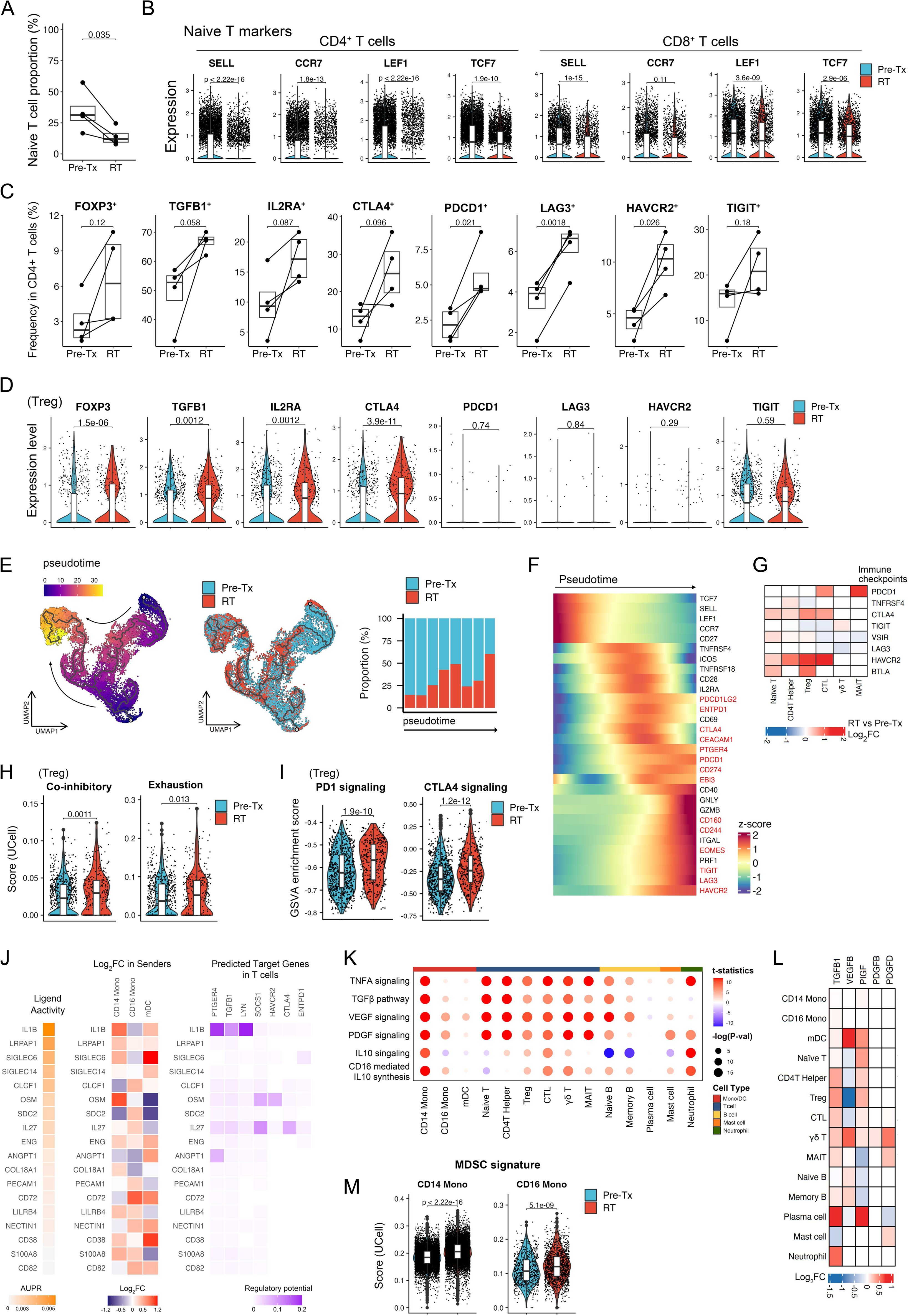
RT promotes progression of immune suppression. (**A)** Proportion of naïve T cells before and after RT. Each line connects paired pre-treatment and post-RT samples from NPC patients. *P*-values were calculated using the paired t test. (**B)** Expression of naïve T cell markers (*SELL*, *CCR7*, *LEF1*, *TCF7*) in CD4⁺ and CD8⁺ T cells pre-treatment (blue) and post RT (red; Mann-Whitney U test). (**C)** Frequency of CD4⁺ T cells expressing Treg-associated markers before and after RT (paired t-test). (**D)** Expression of Treg markers before and after RT (Mann-Whitney U test). (**E)** Pseudotime trajectory analysis of T cells before and after RT. Changes in T cell proportion along pseudotime (right) are shown. (**F**) Changes in the expression of key genes associated with T cell function along pseudotime. Color scale indicates normalized expression levels represented as z-scores, where higher (red) and lower (blue) values reflect relative upregulation or downregulation compared to the gene’s mean expression. (**G)** Log₂ fold changes (log₂FC, RT vs. Pre-Tx) in expression of immune checkpoint genes across T cell subsets. (**H)** Co-inhibitory and exhaustion signature scores in Tregs before and after RT, calculated using the UCell R package (Mann-Whitney U test). (**I)** GSVA analysis of PD-1 and CTLA-4 signaling in Tregs before and after RT. (**J)** Ligand-receptor interaction analysis by NicheNet. Orange, higher AUPRC values (ligand activity); red, upregulated DEGs (log₂FC) in sender cells; blue, downregulated DEGs in sender cells; purple (varying intensity), predicted regulatory target genes in T cells. (**K)** Extrinsic immunosuppressive pathway activity across immune cell types. Dot plots show t-statistics (color) and −log (*P*-value) (dot size). (**L)** Changes in the expression (log₂FC) of extrinsic immunosuppressive genes (*TGFΒ1*, *VEGFB*, *PIGF*, *PDGFB*, *PDGFD*) across immune cell types. (**M)** MDSC signature scores for CD14⁺ and CD16⁺ monocytes, calculated using the UCell R package (Mann-Whitney U test).

A pseudotime trajectory analysis showed a distinct shift in T cell differentiation after RT, with UMAP projections displaying divergent trajectories compared with pre-treatment samples (figure 3E). The T cell proportion increased along pseudotime in post-RT samples (figure 3E, right). The corresponding heatmap showed a gradual decrease in naïve T cell marker expression and upregulation of immunosuppressive genes (figure 3F, marked in red), suggesting that RT promotes a suppressive transcriptional profile in T cells. Expression of inhibitory checkpoint molecules, including *PDCD1* (PD-1), *CTLA4*, *LAG3*, and *HAVCR2*, was significantly upregulated across various T cell subsets post RT (figure 3G). Further characterization of Tregs showed a significant increase in co-inhibitory and exhaustion signature scores post RT (figure 3H), reinforcing the idea that RT enhances an immunosuppressive shift. A GSVA analysis confirmed that RT significantly upregulated PD-1 and CTLA4 signaling pathways in Tregs (figure 3I).

A ligand-receptor interaction analysis using NicheNet identified IL-1β, OSM (oncostatin M), and IL-27 as key mediators of T cell immune regulation following RT (figure 3J). These cytokines, primarily derived from myeloid and stromal cells, exhibited increased ligand activity post RT, as indicated by higher AUPRC (area under the precision-recall curve) scores. A differential expression analysis confirmed upregulation of these cytokines in CD14⁺ monocytes, CD16⁺ monocytes and mDCs, suggesting RT-induced reprogramming of the tumor immune microenvironment. Predicted target genes in T cells, including *PTGER4*, *TGFB1*, *LYN* (Src family tyrosine kinase), *SOCS1* (suppressor of cytokine signaling 1), *HAVCR2*, *CTLA4* and *ENTPD1* (ectonucleoside triphosphate diphosphohydrolase 1) were enriched and regulatory functions governed by the corresponding proteins were upregulated, supporting a role for these interactions in driving Treg-mediated immunosuppression.

In addition to direct T cell modulation, RT also increased the activity of signaling pathways associated with the extrinsic immunosuppressive factors, TGFβ1, VEGFB, PIGF (phosphatidylinositol glycan anchor biosynthesis class F), PDGFB (platelet-derived growth factor A) and PDGFD, across multiple immune cell types (figure 3K and L, and online supplemental Table 3). Notably, RT enhanced the MDSC signature in CD14⁺ and CD16⁺ monocytes (figure 3M). Collectively, these findings suggest that RT promotes both Treg-driven and broader immunosuppressive signaling in the TME.

### RT combined with the STING agonist diABZI enhances therapeutic efficacy and modulates immune responses

Our scRNA-seq analysis revealed that fractionated RT suppresses the cGAS–STING pathway and enhances the immunosuppressive Treg signature. This potential causal relationship prompted us to test whether a STING agonist could counteract RT-induced immune suppression. To this end, we evaluated the effect of the STING agonist, diABZI, alone or in combination with fractionated RT (12 Gy in 4-Gy fractions), in a syngeneic mouse tumor model (figure 4A). Photon therapy more effectively suppressed tumor growth than electron beam therapy; however, neither modality had fully controlled tumor progression by the end of the observation period (figure 4B and C). diABZI significantly enhanced tumor control both as a monotherapy and in combination with RT (figure 4D and E, and online supplemental Table 4). Mice treated with diABZI alone or in combination with RT exhibited enlarged spleens (figure 4 F). The RT-induced immune response and the STING agonist-induced effects on mouse spleens and tumor tissues were dissected by analyzing bulk RNA sequencing data. Importantly, these in vivo results confirmed that RT induces suppression of cGAS–STING signaling (figure 4F). A Gene Set Enrichment Analysis (GSEA) revealed the functions most affected by RT and diABZI. Notably, diABZI stimulation enhanced neutrophil degranulation and inhibited PD-1 signaling in both spleens and tumor tissues of mice treated with diABZI alone or in combination with RT (figure 4G-H), reversing the expression of virtually all PD-1 signaling signature genes in both tumors and spleens (figure 4I). These results identify mechanisms underlying RT- and STING agonist-induced immune modulation in systemic immunity as well as tumor immunity.

**Figure 4.**
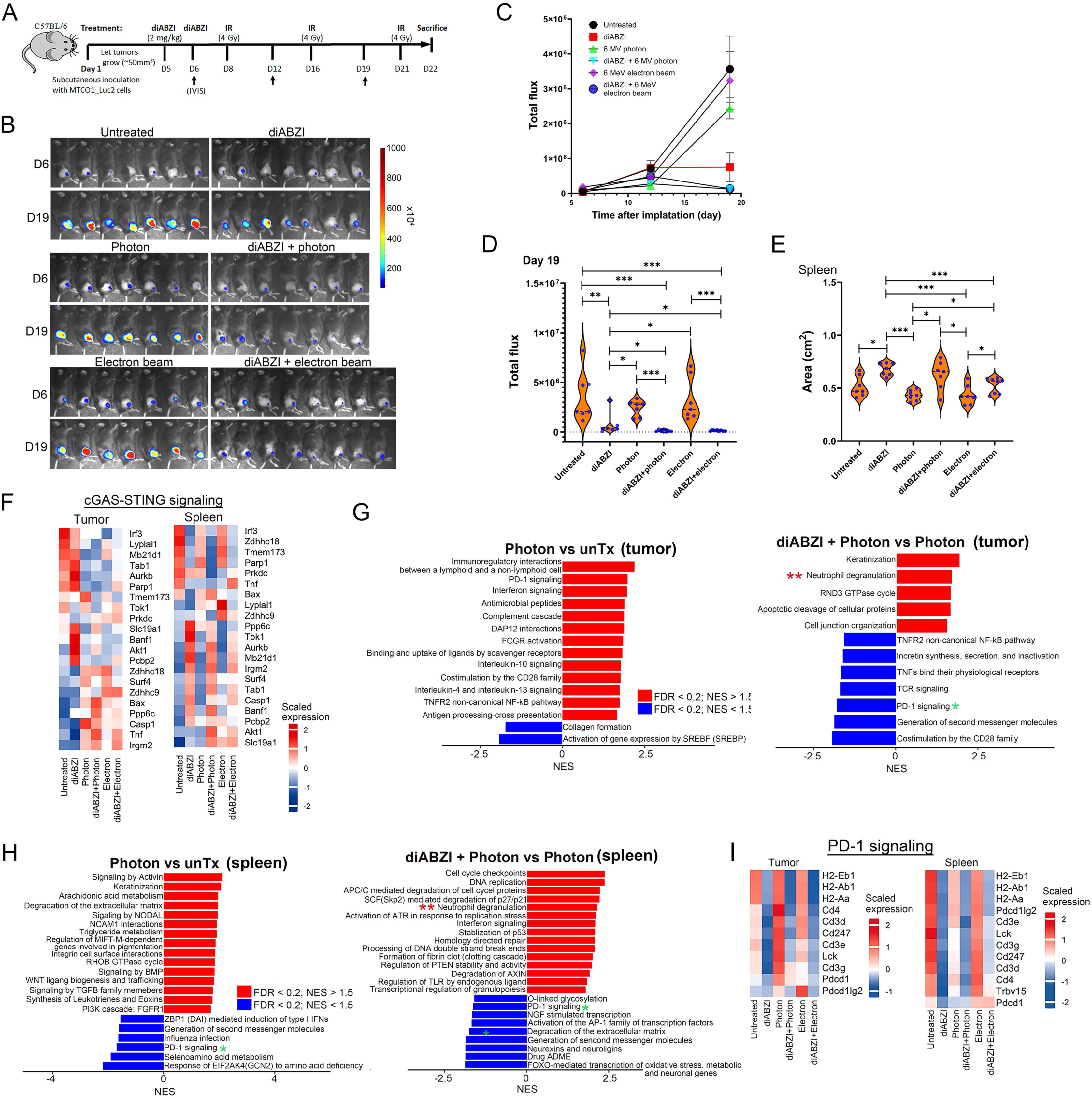
RT combined with the STING agonist diABZI enhances therapeutic efficacy and modulates immune responses. (**A)** Schematic of the treatment protocol in mouse MTCQ1_Luc2 tumor-bearing C57BL/6 mice treated with diABZI and/or RT. Day 1 indicates the time of tumor cell inoculation. diABZI (2 mg/kg, intraperitoneal) and local tumor RT (6 MV photon or 6 MeV electron beam) were administered as indicated. Tumor progression was monitored using an in vivo bioluminescence imaging system (IVIS). (**B)** IVIS images on days 6, and 19 for the indicated treatment groups: untreated, diABZI alone, photon irradiation alone, diABZI + photon irradiation, electron beam alone, and diABZI + electron beam. (**C)** Tumor growth curves showing total flux (photons/s) from IVIS measurements. Data are presented as means ± SD. (**D)** Tumor burden on day 19 based on total flux. Each dot represents one mouse (n = 7– 8 mice/group; \**P* < 0.05, \*\**P* < 0.01, \*\*\**P* < 0.001; Mann–Whitney test). (**E**) Quantification of spleen area (\**P* < 0.05, \*\*\**P* < 0.001; Mann–Whitney test). (**F)** Heatmaps of cGAS–STING signaling genes in tumors and spleens (P < 0.05, |log2FC| > 1). Data are presented as means of replicates (n = 2 for untreated, n = 3 or 4 for other groups). Red, upregulation; blue, downregulation. (**G**) GSEA in tumors comparing photon versus untreated (left) and diABZI + photon versus photon alone (right). (**H**) GSEA in spleens comparing photon versus untreated (left) and diABZI + photon versus photon alone (right). Red, increased normalized enrichment score (NES); blue, decreased NES. (**I**) Heatmaps of PD-1 pathway genes in tumors and spleens. Data are presented as means of replicates. Red, upregulation; blue, downregulation.

### diABZI stimulation enhances neutrophil activation in association with improved tumor control

To further characterize the biological relevance of diABZI-induced neutrophil degranulation with therapeutic outcomes, we analyzed alterations in genes involved in neutrophil degranulation among groups in our in vivo model. Results of transcriptomic analyses showed that, overall, diABZI treatment alone or in combination with RT increased the neutrophil-degranulation expression signature, particularly in spleens (figure 5A). Myeloperoxidase (MPO) plays crucial role in modulating neutrophil functions [38], but the association between MPO expression and tumor progression is context-dependent [39–41]. Immunohistochemical results showed that diABZI increased the abundance of MPO-positive neutrophils in the spleen (figure 5B and C), suggesting enhanced neutrophil activation. Moreover, MPO levels were positively correlated with spleen size (r = 0.4128, *P* = 0.0059; figure 5D). MPO protein abundance in tumor tissues was markedly increased in the diABZI + electron group, and moderately increased in the diABZI + photon group and the electron treatment-only group (figure 5E and F). Importantly, spleen MPO expression inversely correlated with tumor burden (r-= −0.7038, *P* < 0.0001; figure 5G), suggesting that enhanced innate immune activity is associated with better tumor control.

**Figure 5.**
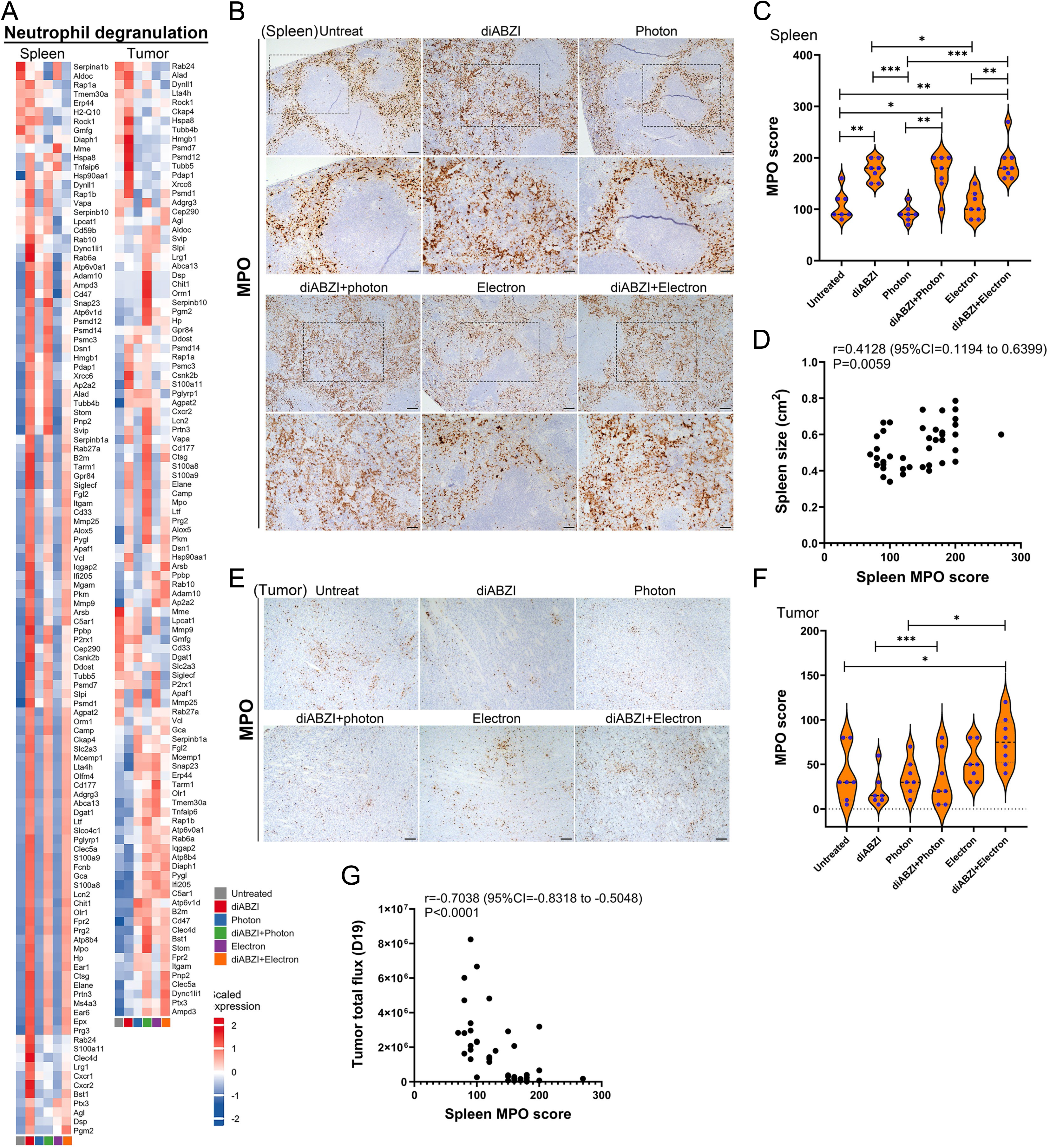
Activation of neutrophil-degranulation responses following diABZI treatment. (**A**) Heatmaps of neutrophil degranulation-related genes in spleens (left) and tumors (right). Data are presented as means of replicates. Red, upregulation; blue, downregulation. (**B**) Representative IHC images of MPO expression in spleen sections. Scale bars, 100 μm (low magnification) and 50 μm (high magnification. (**C**) Quantification of MPO scores in spleens (\**P* < 0.05, \*\**P* < 0.01, \*\*\**P* < 0.001; Fisher’s exact test). (**D**) Correlation between spleen MPO levels and spleen size (Spearman’s correlation test). (**E**) Representative IHC images of MPO in tumors. Scale bars, 50 μm. (**F**) Quantification of tumor MPO scores (\**P* < 0.05, \*\*\**P* < 0.001; Fisher’s exact test). (**G**) Correlation between spleen MPO scores and tumor burden on day 19 (Spearman’s correlation test).

### ATM signaling plays a crucial role in RT-induced immune suppression

Signaling mediated by the serine/threonine kinase, ATM, regulates crucial cell functions, including cell-cycle checkpoints, DNA repair, and innate and adaptive immunity, in response to DNA damage and reactive oxygen species (ROS) [42–44]. Our transcriptomic analysis revealed that a majority of genes involved in ATM signaling were downregulated in the spleens of untreated and RT groups, but showed an enhanced ATM signature in the spleens of diABZI-treated groups (figure 6A, left), indicating the involvement of ATM signaling in STING-mediated immune activation. In tumors, RT suppressed the expression of ATM signature genes, an effect that was partly reversed by combined treatment with diABZI (figure 6A, right). Immunohistochemical (IHC) examination of activated ATM (S1981) in spleens revealed a dramatic diABZI-induced enhancement of phosphorylated ATM (p-ATM) (figure 6B and C). In tumor tissues, RT induced activation of ATM compared to that in the untreated group, and diABZI synergistically enhanced RT-induced ATM activation (figure 6D). Importantly, spleen p-ATM levels were inversely correlated with tumor burden (r = −0.6696, *P* < 0.0001; figure 6E). A weaker, but significant, inverse correlation was also observed between tumor p-ATM levels and tumor burden (r = −0.3256, *P* = 0.0331; figure 6F). Further correlation analyses revealed a positive association between p-ATM (S1981) and MPO IHC scores in both spleens (r = 0.7013, *P* < 0.0001; figure 6G) and tumor tissues (r = 0.4349, *P* = 0.0036; figure 6H). Additionally, elevated ATM levels in NPC tumors (GSE102349) were significantly associated with better progression-free survival compared to tumors with low ATM expression (*P* = 0.021, log-rank test; figure. 6I). Moreover, cGAS–STING and ATM signaling were highly correlated in NPC tissues (r = 0.7, P < 2.2 × 10⁻¹⁶; figure 6J). To further confirm that diABZI indeed regulates ATM signaling, MPO and the innate immune response, we assessed the expression of key molecules involved in these functions in various cell types, including PBMCs, the THP-1 cell line (monocytes), THP-1–derived macrophages, and the Jurkat cell line (CD4 Th cells). This analysis showed that various innate immune proteins, including S366-phosphoryated STING (p-STING), S396-phosphorylated interferon regulator factor (p-IRF), CASP1, PARP (poly(ADP-ribose) polymerase) and IL-1β, in diABZI-treated cells, were activated to varying degrees and for different durations (figure 6K–N). diABZI induced upregulation of MPO in PBMCs and THP-1 monocytes at 3 and 6 hours post treatment (figure 6K and 6M), promoted ATM activation in PBMCs and THP-1 monocytes, and activated γH2AX (S139) in all immune cell lines tested. Further, with the exception of THP-1-derived macrophages, diABZI enhanced ATM activation in cells exposed to cisplatin-induced DNA damage, suggesting that diABZI activates both innate immune and DNA-damage responses. Together, these findings suggest that ATM signaling plays a crucial role in modulating systemic immune responses and tumor control, and that ATM activation is associated with increased neutrophil infiltration (MPO expression), and that STING-ATM activation is associated with enhanced anti-tumor immunity.

**Figure 6.**
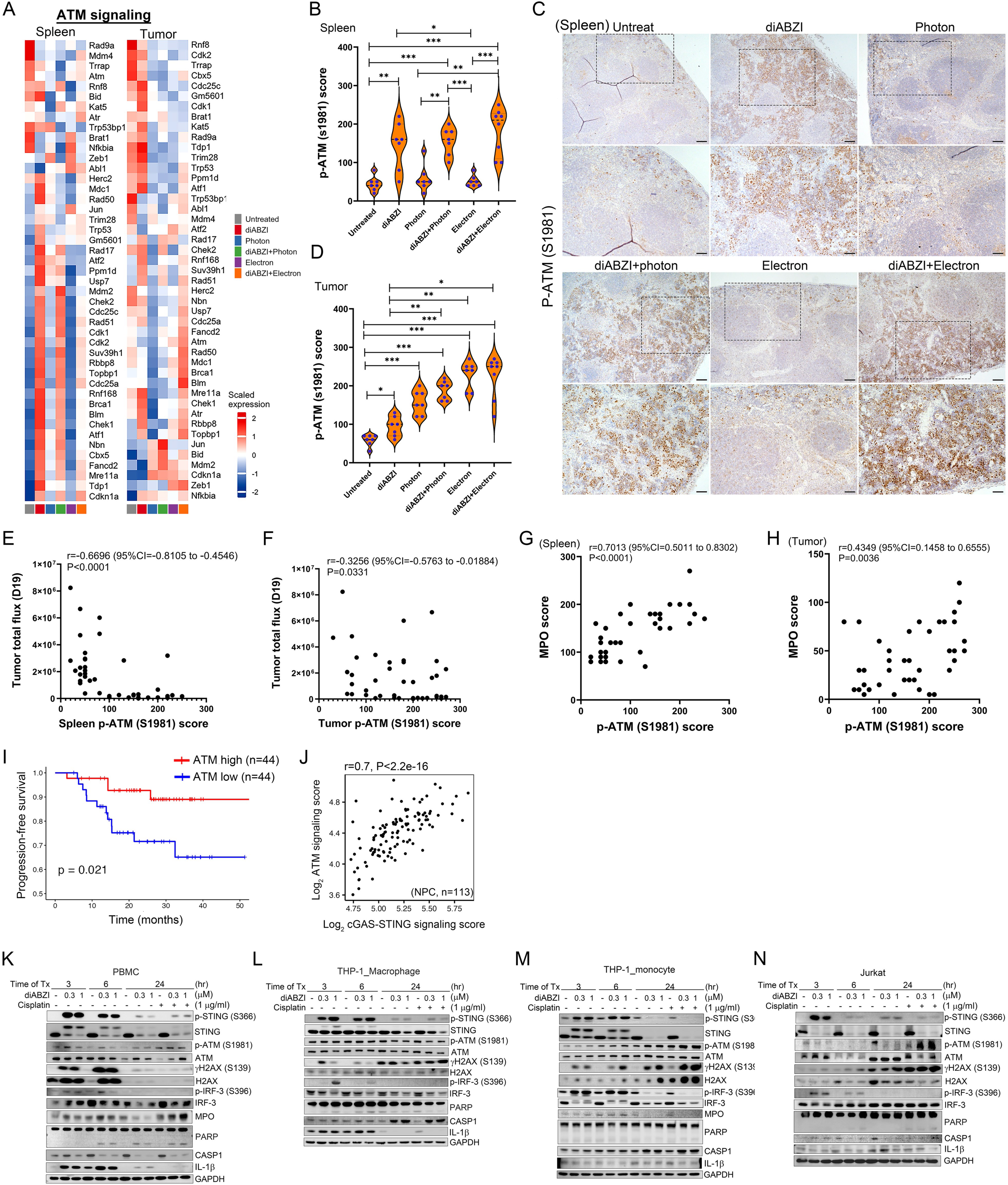
ATM signaling mediates the therapeutic effect of diABZI. (**A**) Heatmaps of ATM signaling genes in spleens (left) and tumors (right). Data are presented as means of replicates. Red, upregulation; blue, downregulation. (**B)** Quantification of p-ATM (S1981) scores in spleens across treatment groups (\**P* < 0.05, \*\**P* < 0.01, \*\*\**P* < 0.001; Fisher’s exact test). **(C)** Representative IHC images of p-ATM (S1981) in spleens. Scale bars, 100 μm and 50 μm (high-magnification). (**D**) Quantification of p-ATM (S1981) scores in tumors (\**P* < 0.05, \*\**P* < 0.01, \*\*\**P* < 0.001; Fisher’s exact test). (E**F** and **F)** Correlation of p-ATM (S1981) scores and tumor burden in spleens (**E**) and tumors (**F**) on day 19 (Spearman’s correlation test). **(G** and **H**) Correlation between p-ATM (S1981) and MPO scores in spleens (**G**) and tumors (**H**). Spearman’s correlation coefficient (r), 95% confidence interval (CI), and *P*-values are indicated. (**I**) Kaplan Meier progression-free survival curves for NPC patients stratified by ATM expression in tumors (log-rank test). (**J)** Correlation of cGAS–STING with ATM signaling scores in NPC tumors (Spearman’s correlation test). Data for (K) and (L) are extracted from the GEO database (GSE102349). (**K–L)** Western blot analysis of innate immune and DNA damage-response proteins in PBMCs (**K**), THP-1–derived macrophages (**L**), THP-1 monocytes (**M**), and Jurkat cells (**N**) treated with diABZI (0.3 or 1 μM), cisplatin (1 μg/mL), or both for 3, 6, or 24 hours.

### diABZI-induced ATM activation is correlated with reduced Treg abundance and improved tumor control

Transcriptomic data further revealed that fractionated RT enhanced immune coinhibitory signaling and IL-10 signaling in tumors (figure 7A). Results of IHC showed that FOXP3 immunoreactivity was most abundant in untreated tumors (figure 7B and C). Combinational treatment with diABZI + photons resulted in a reduction in FOXP3 immunoreactivity. IHC showed a negative association between activated ATM immunoreactivity and FOXP3 abundance in tumors (figure 7D), and revealed a positive correlation between FOXP3 expression and TGFB1 expression (figure 7E). Notably, a correlation analysis revealed a significant inverse relationship between p-ATM and FOXP3 expression in tumor tissues (r = −0.778, *P* < 0.0001; figure 7F). We performed double immunofluorescence staining for p-ATM (S1981) and FOXP3 in tissues from untreated and diABZI + electron beam-treated groups, representing the most immunosuppressive and activated immune environments, respectively. diABZI + electron beam treatment induced numerous p-ATM foci (green), whereas FOXP3 signals were rarely detected in tumors (figure 7G) or spleens (figure 7H) compared with untreated controls. Conversely, clusters of FOXP3^+^ immune cells (red) were observed in tissues with low p-ATM immunoreactivity. These data suggest that diABZI-induced ATM activation is associated with reduced Treg expansion.

**Figure 7.**
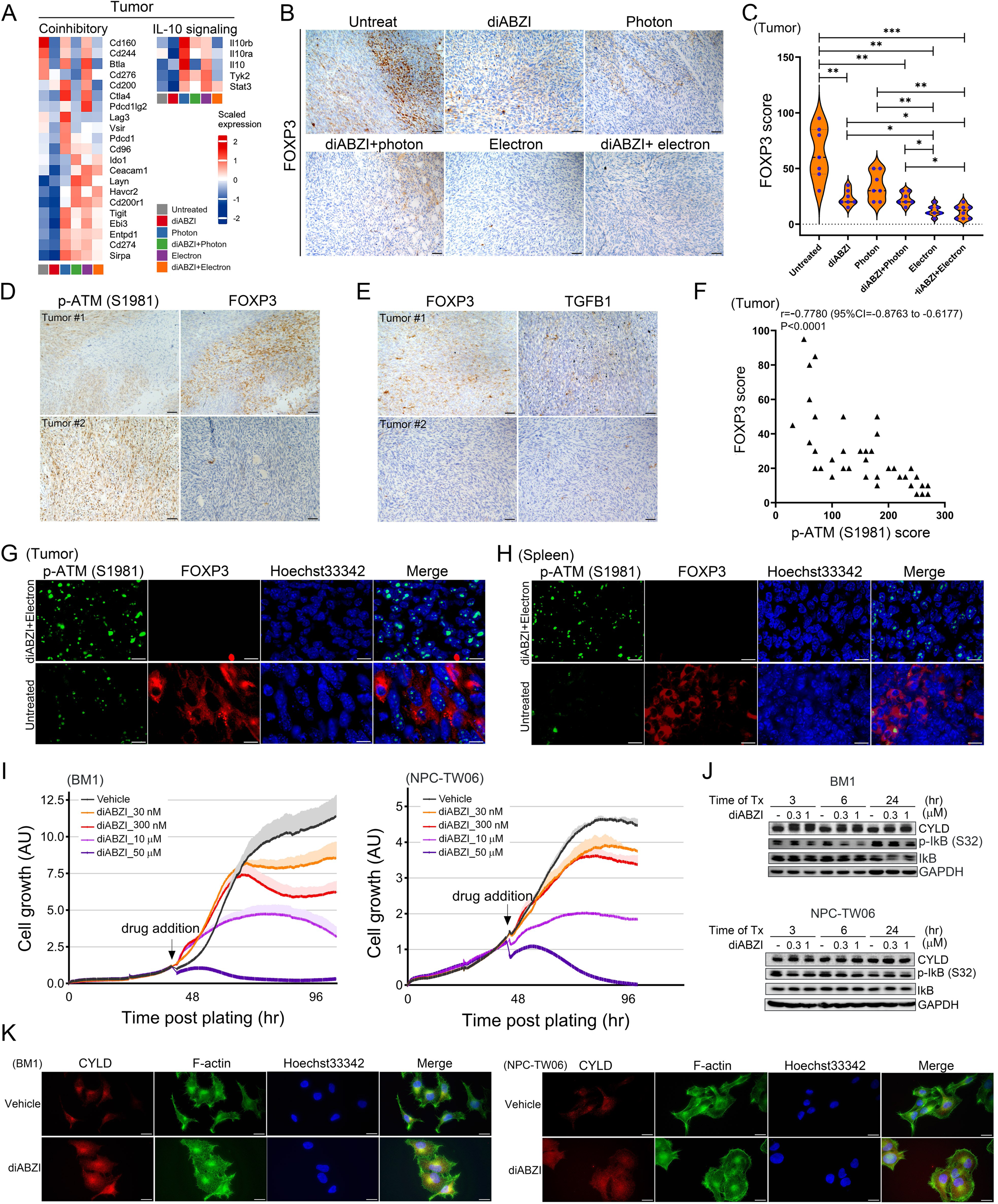
diABZI treatment reduces immune suppression and Treg expansion. (**A**) Heatmaps show coinhibitory and IL-10 signaling signatures in tumor tissues from the indicated groups. Data are presented as means of replicates. Red, upregulation; blue, downregulation. (**B)** Quantification of FOXP3 scores in tumors across groups (\**P* < 0.05, \*\**P* < 0.01, \*\*\**P* < 0.001; Fisher’s exact test). (**C)** Representative IHC images of FOXP3 expression in tumors across all groups. Scale bars, 50 μm. (**D**) Representative IHC images of p-ATM and FOXP3 expression in consecutive tumor sections. Scale bars, 50 μm. (**E)** Representative IHC images of FOXP3 and TGFB1 expression in consecutive tumor sections. Scale bars, 50 μm. (**F)** Correlation analysis between p-ATM (S1981) and FOXP3 expression in tumors (Spearman’s correlation test). (**G** and **H**) Immunofluorescence staining for p-ATM (S1984) (green) and FOXP3 (red) in tumors (**G**) and spleens (**H**) from untreated and diABZI + electron beam-treated groups. (**I**) Real-time monitoring of NPC cell growth in response to diABZI stimulation using an xCelligence real-time analyzer. Values represent means and SD of triplicate experiments. Black arrows indicate the time of diABZI addition. (**J**) Western blot analysis of CYLD and phosphorylated IκB (S32) levels in diABZI-treated NPC cells. GAPDH was used as a loading control. (**K**) Immunofluorescence staining of CYLD in NPC cells treated with diABZI (0.3 μM) for 3 hours. F-actin was stained with Alexa Fluor 488 phalloidin (green), and nuclei were counterstained with Hoechst33342 (blue). Scale bars, 20 μm.

We further assessed the effects of the STING agonist on NPC cells and found that, similar to immune populations, diABZI treatment activated STING signaling and DNA damage response molecules. However, it also induced morphological abnormalities and cell death after 24 hours in a concentration-dependent manner, while vehicle-treated cells continued proliferating, showing mitotic activity even under low-serum conditions. Moreover, combined treatment with diABZI and cisplatin led to increased cell death compared to treatment with either diABZI or cisplatin alone. To quantify cytotoxicity, we treated NPC cells with vehicle (DMSO) or varying doses of diABZI (30 nM, 300 nM, 10 μM, and 50 μM), and monitored cell growth using an xCelligence real-time cell analyzer. As shown in figure 7I, diABZI induced dose-dependent cytotoxicity. Western blot analysis revealed that diABZI stimulation increased levels of cylindromatosis (CYLD), a known negative regulator of NF-κB [45], and reduced phosphorylation of IκB (Ser 32) in diABZI-treated NPC cells (figure 7J). In addition, CYLD expression was enhanced in diABZI-treated cells, in contrast to the low cytosolic expression observed in controls (figure 7K). Taken together, these results suggest that diABZI treatment not only activates anti-tumor immunity but also directly induces cytotoxic effects in NPC cells, potentially through CYLD upregulation and NF-κB pathway inhibition, thereby contributing to tumor regression.

Collectively, our findings establish a mechanistic link between RT-induced immune suppression and inhibition of cGAS–STING signaling in NPC, and show that activation of the STING–ATM–MPO axis using a STING agonist restores anti-tumor immunity, reduces Treg expansion and improves tumor control, supporting the therapeutic potential of STING agonists in combination with RT against NPC.

## Discussion

In this study, we analyzed the systemic immune responses to radiotherapy (RT) in a rare cohort of stage I (T1N0M0) NPC patients treated with RT alone. This unique design allowed us to isolate RT-specific effects without the confounding influence of concurrent chemotherapy. Notably, our scRNA-seq analysis of paired PBMCs revealed that fractionated RT suppressed cGAS–STING signaling in monocytes and dendritic cells, despite an overall increase in DNA damage responses. This suggests selective impairment of cytosolic DNA sensing post-irradiation. RT also induced an immunosuppressive shift in T cells, including enhanced expression of *FOXP3*, *CTLA4*, and *PDCD1*, and a reduction in naïve markers, indicative of Treg expansion and T cell exhaustion.

Given the systemic nature of RT-induced immune modulation and the known abscopal effects of RT, we focused on circulating immune populations. Tumor sampling during therapy remains clinically impractical, further justifying this approach. Moreover, systemic immune monitoring provides insight into therapeutic resistance mechanisms and opportunities for immunomodulation.

Previous preclinical studies have shown that STING activation can enhance RT efficacy in mouse models of pancreatic cancer [46] and glioblastoma [47]. Our study expands on this by incorporating several key differences: (1) we evaluated RT-induced immune remodeling in a human clinical cohort receiving RT alone, enabling clear attribution of observed changes to RT; (2) in the syngeneic mouse tumor model, we used a clinically relevant, fractionated RT regimen instead of single high-dose irradiation; and (3) we compared immune responses under photon and electron beam irradiation with equivalent dosing, revealing differential effects on downstream signaling and immune composition.

To counteract RT-induced immune suppression, we investigated the STING agonist diABZI. In vivo, the combination of RT and diABZI restored STING and ATM signaling, enhanced neutrophil degranulation, suppressed PD-1 signaling and Treg expansion, and improved tumor control. Notably, electron beam treatment, either alone or combined with diABZI, resulted in the highest levels of MPO and p-ATM and the lowest FOXP3 abundance in tumors. This may be attributable to the direct ionization capacity of electron beams, although their limited penetration restricts their clinical applicability in deep-seated tumors such as NPC.

In addition to its immunomodulatory effects, diABZI directly exerted cytotoxic activity against NPC cells. In vitro assays demonstrated dose-dependent growth inhibition, nuclear translocation of IRF3, and CYLD-mediated suppression of NF-κB signaling, leading to increased cell death. These findings challenge the previous notion that STING agonists lack direct anti-tumor effects in NPC and underscore their dual function as immune activators and tumoricidal agents.

Despite these promising results, several translational challenges remain. STING agonists face limitations in pharmacokinetics, delivery, and toxicity. Recent advances in nanoparticle-based delivery systems [48], such as NP-cGAMP for inhalable administration in a mouse lung metastasis tumor model [49] and NAcp@CD47 for targeted dual delivery in glioblastoma tissues [50], provide a promising strategy to improve therapeutic efficacy and tissue specificity. Nonetheless, the long-term safety, optimal delivery modalities, and combinatorial strategies with RT, chemotherapy, or immune checkpoint blockade remain to be fully explored.

Lastly, RT delivery parameters may critically influence therapeutic outcome. High-dose RT can induce TREX1 expression, degrading cytosolic DNA and suppressing cGAS–STING activation [20]. Optimizing fractionation schedules to preserve DNA-sensing pathways may be essential for maximizing synergy with STING agonists.

## Acknowledgments

We thank Ms. Hsing-Ju Cheng and Mr. Yi Huang from the Department of Radiation Oncology, Taipei Tzu Chi Hospital, for performing mouse tumor irradiation. We appreciate the support from the Bioinformatics Core of the National Health Research Institutes, funded by the National Core Facility for Biopharmaceuticals (NCFB), National Science and Technology Council (NSTC113-2740-B-400-005). We would like to thank National Core Facility for Biopharmaceuticals (NCFB, NSTC 114-2740-B-492-001) and National Center for High-performance Computing (NCHC) of National Institutes of Applied Research (NIAR) of Taiwan for providing computational and storage resources.

## Ethics approval

Ethical approval was granted by the Institutional Review Board of Chang Gung Memorial Hospital, Taiwan (IRB numbers: 202000190B0, 202101548B0). All human specimens were handled in accordance with institutional and national guidelines. Animal experiments were conducted in compliance with the principles of laboratory animal care and were approved by the animal committees of Chang Gung University (CGU110-161) and National Central University (NCU-112-026).

